# Lineage-restricted dependency on an oncofetal *SNHG29*-IGF2BP1 RNA axis in acute megakaryoblastic leukemia

**DOI:** 10.64898/2026.02.07.704501

**Authors:** Robert Winkler, Bruno Griesler, Wolfgang Sippl, Reinier A. Boon, Ilka Wittig, Marie-Laure Yaspo, Stefan Hüttelmaier, Raj Bhayadia, Dirk Heckl, Jan-Henning Klusmann

## Abstract

Acute megakaryoblastic leukemia (AMKL) is a rare, aggressive subtype of acute myeloid leukemia with developmental origins in early childhood. To uncover long noncoding RNAs (lncRNAs) sustaining this high-risk malignancy, we conducted CRISPR interference screens targeting lncRNAs overexpressed in primary AMKL samples. This analysis identified *SNHG29* as a previously unrecognized lineage-specific dependency, whose silencing profoundly impaired leukemic proliferation and clonogenic growth in vitro and reduced leukemic burden in vivo. RNA pulldown coupled with proteomic analysis revealed that *SNHG29* interacts with the oncofetal RNA-binding protein IGF2BP1, which is aberrantly expressed in AMKL. *SNHG29* was required to maintain expression of IGF2BP1 target transcripts, including MYC- and E2F-driven proliferative programs, thereby reinforcing fetal transcriptional programs essential for leukemic maintenance. Pharmacologic inhibition of IGF2BP1-RNA interactions with the small-molecule BTYNB induced potent and selective cytotoxicity in patient-derived AMKL models. Together, these findings uncover a developmental co-dependency between *SNHG29* and IGF2BP1 that defines a lineage-restricted oncogenic circuit and an actionable therapeutic vulnerability in AMKL.

## Introduction

Long non-coding RNAs (lncRNAs) are transcripts longer than 200 nucleotides that lack protein-coding potential,^1,2^ but exert diverse regulatory functions. In contrast to protein-coding genes, lncRNAs display highly specific expression patterns, often restricted to distinct cell lineages, developmental stages, or disease contexts, suggesting specialized biological roles.^3,4^ They act through multiple mechanisms – by scaffolding protein complexes^5^, guiding chromatin modifiers^6^, modulating mRNA stability^7^, or sequestering microRNAs^8,9^ – to fine-tune transcriptional and post-transcriptional programs. Recent work has uncovered central roles for lncRNAs in hematopoiesis, where they control hematopoietic stem cell maintenance, lineage commitment, and terminal differentiation^10,11^. Dysregulated lncRNA expression contributes to malignant transformation across hematologic neoplasms,^12–14^ including acute leukemias, underscoring their potential as therapeutic targets.

Acute megakaryoblastic leukemia (AMKL) is a rare and aggressive subtype of acute myeloid leukemia (AML) characterized by uncontrolled proliferation of megakaryoblasts. AMKL has developmental ties to early hematopoiesis and frequently arises in early childhood^15^, particularly in the context of Down syndrome (ML-DS), which has a favorable outcome.^16,17^ In contrast, non–Down syndrome AMKL (non-DS-AMKL) is driven by high-risk fusion oncogenes such as *CBFA2T3::GLIS2*, *NUP98::KDM5A* or *RBM15::MKL1*,^18,19^ and is associated with poor survival despite intensified chemotherapy,^20,21^ highlighting the urgent need for innovative, molecularly guided therapies.

We hypothesized that the distinct transcriptional landscape of AMKL – shaped by developmental lineage programs and fusion-driven oncogenic signaling – creates dependencies on specific long non-coding RNAs that sustain leukemic identity and survival through post-transcriptional regulatory mechanisms. To test this hypothesis, we performed a high-throughput CRISPR interference (CRISPRi) screen targeting lncRNAs that are overexpressed in primary AMKL samples. This approach identified *SNHG29* as a selective and essential dependency. Using complementary genetic perturbation strategies, we established its functional relevance in vitro and in vivo, delineated its protein interactome, and uncovered a mechanistic link to the oncofetal RNA-binding protein (RBP) IGF2BP1. We further explored therapeutic disruption of this lncRNA–RBP axis. Together, our findings identify *SNHG29* as a critical lncRNA in AMKL and reveal a previously unrecognized *SNHG29*–IGF2BP1 regulatory axis that represents an actionable vulnerability in megakaryoblastic leukemia.

## Results

### Systematic Investigation of AMKL lncRNAs using CRISPRi Screens

To define the transcriptional landscape underlying AMKL, we performed comprehensive lncRNA expression profiling across FACS-sorted hematopoietic stem cells, progenitors, mature blood lineages, and pediatric AML blasts spanning multiple cytogenetic subtypes. This analysis revealed a distinct lncRNA signature selectively enriched in AMKL blasts relative to all other hematopoietic populations and AML subtypes (**Figure 1A**).^22^

**Fig. 1.**
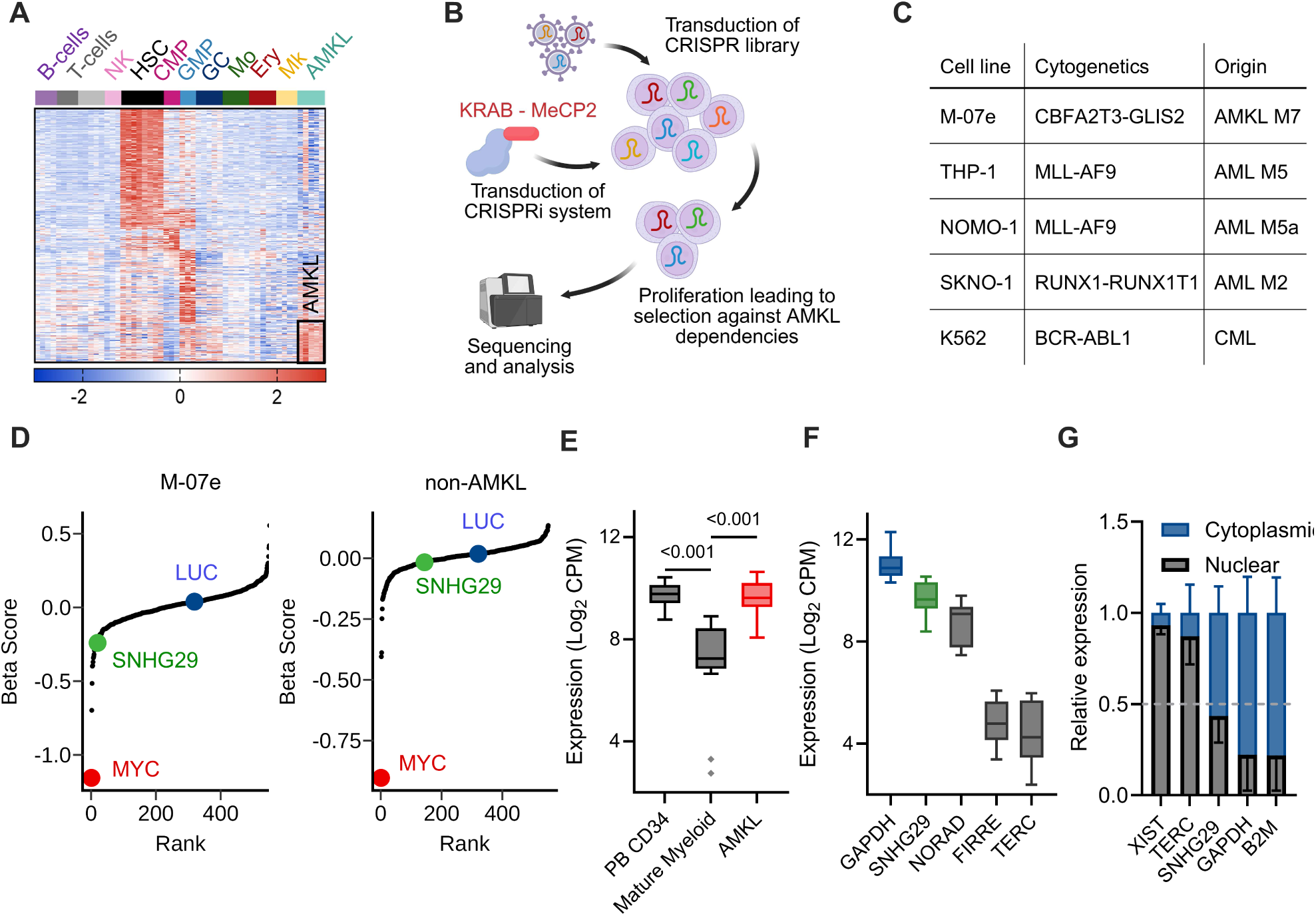
**A** LncRNA expression profiling in primary AMKL patient samples and a panel of normal hematopoietic cell populations. Differential analysis reveals a unique signature enriched in AMKL, which is highlighted by the black rectangle. expression values are shown as row-wise z-scores. NK: Natural killer cell; HSC: Hematopoietic stem cell; CMP: Common myeloid progenitor; GMP: Granulocyte-monocyte progenitor; GC, Granulocyte; Mo, Monocyte; Ery, Erythroid precursor; Mk, Megakaryocyte; AMKL, Acute megakaryoblastic leukemia patient samples. **B** Schematic of the CRISPRi screening workflow designed to functionally interrogate the AMKL lncRNA signature. **C** Cell lines included in the screening, their cytogenetic characteristics and disease of origin. **D** CRISPRi screen results showing gene essentiality scores in M-07e cells (left) and all other cell lines (right), calculated by MAGeCK MLE. Key hit *SNHG29*, positive control *MYC* and negative control *LUC* are highlighted in pink, green and violet, respectively. **E** Expression levels of *SNHG29* in RNA sequencing datasets from peripheral blood mobilized CD34+ cells (PB CD34, n=8), differentiated myeloid cells (n=16, monocytes, late megakaryocytic and erythroid progenitors, granulocytes), and AMKL patient samples (n=17). Data shown as LIMMA-voom transformed counts per million on Log2 scale. ****FDR<0.0001 (LIMMA-voom). Box plots show medians, boxes and whiskers according to the Tukey method. **F** Expression levels of *SNHG29* (pink), other lncRNAs (green) and protein-coding genes (grey) in AMKL patient samples (n=17). Data shown as LIMMA-voom transformed counts per million on g2 scale. Box plots show medians, boxes and whiskers according to the Tukey method. **G** *SNHG29* abundance in nuclear (grey) versus cytoplasmic (pink) fractions, quantified by qRT-PCR. Localization is compared to established nuclear (XIST, TERC) and cytoplasmic (GAPDH, B2M) control transcripts. Data are presented as percentage of total detected transcript in each fraction (n=2, mean ± s.e.m.).

To assess the functional relevance of these lncRNAs, we performed a high-throughput CRISPR interference (CRISPRi, dcas9.KRAB.MeCP2)^23^ screen targeting 548 noncoding RNA genes (**Figure 1B**). The screen was carried out in the AMKL-derived M-07e line, which harbors a *CBFA2T3::GLIS2* fusion, alongside a reference panel of AML cell lines representing alternative subtypes (**Figure 1C**). Among the top-scoring hits, *SNHG29* emerged as a strong and selective dependency in AMKL, with its knockdown markedly reducing M-07e viability (**Figure 1D**), while having minimal impact on non-AMKL cell lines. Consistently, *SNHG29* was highly expressed in AMKL patient samples and downregulated during normal hematopoietic differentiation (**Figure 1E**), is abundant relative to other lncRNAs (**Figure 1F**) and exhibits both nuclear and cytoplasmic localization (**Figure 1G**). Together, these results identify *SNHG29* as a candidate lineage-specific dependency lncRNA in AMKL.

### SNHG29 is a lineage-specific dependency lncRNA in AMKL

To validate the CRISPRi screen results, we employed complementary loss-of-function strategies including shRNA-mediated knockdown, RNA-targeting by CRISPR-Cas13d (RfxCas13d, CRISPR-CasRx), and CRISPR-Cas9-mediated genomic deletion (**Figure 2A**). All perturbations confirmed that *SNHG29* is essential for AMKL cell proliferation (**Figure 2B**). shRNA-mediated knockdown of *SNHG29* significantly impaired M-07e growth, with effects proportional to knockdown efficiency (**Figure 2C**). CRISPR-CasRx also recapitulated these results.

**Fig. 2.**
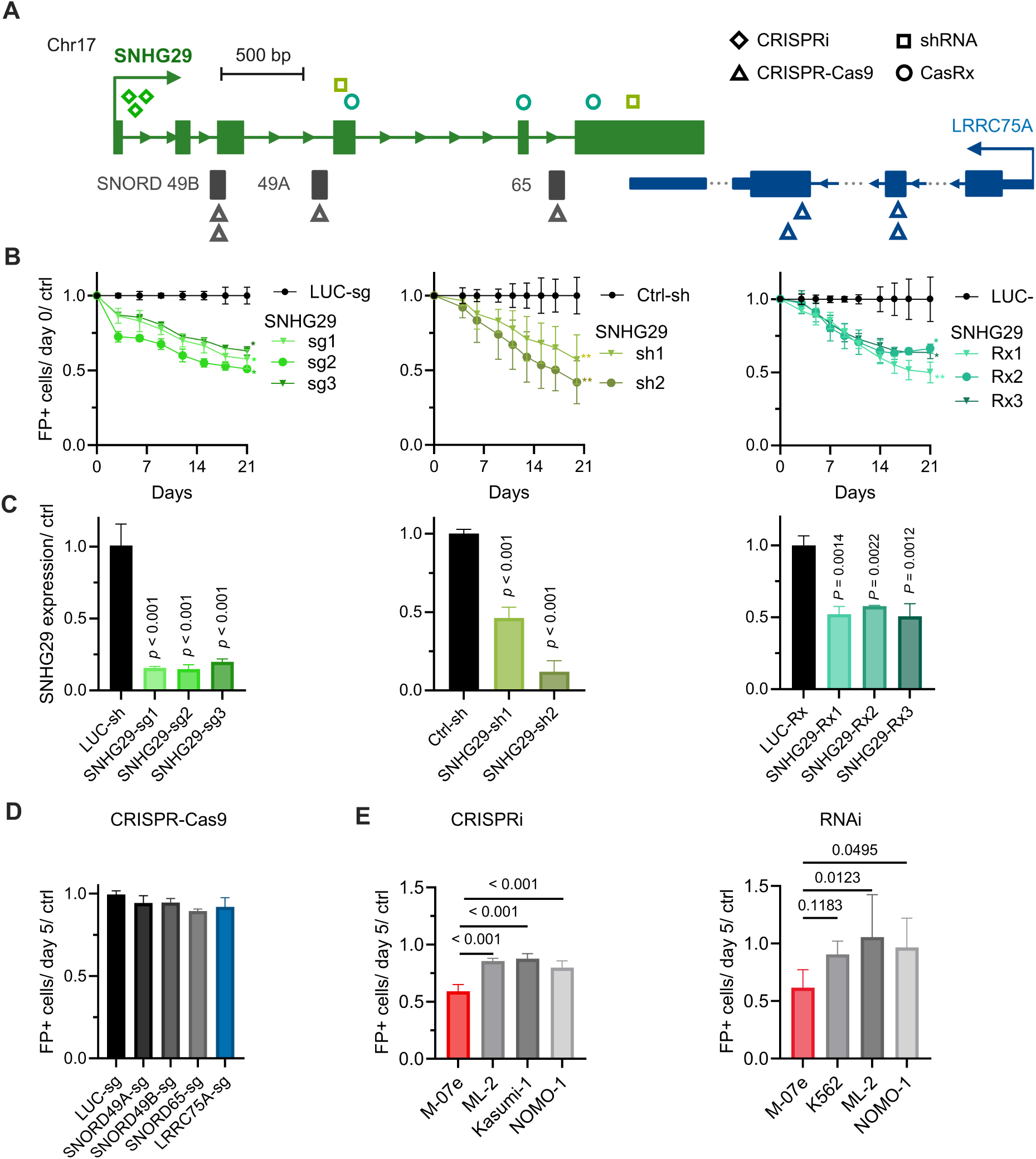
**A** Schematic representation of the *SNHG29* genomic locus on chromosome 17. Arrow indicates transcriptional orientation. Boxes and connecting lines represent exons and introns of *SNHG29*, respectively. Symbols at the bottom indicate target sites for different perturbation strategies: CRISPRi (diamonds), shRNA (squares), CRISPR-CasRx (circles) and CRISPR-Cas9 (triangles). **B** Fluorescence-based proliferation assays in M-07e cells using different *SNHG29* knockdown strategies. Left: CRISPRi using three sgRNAs (n=2). Middle: RNAi using two shRNAs (n=5 for *SNHG29*-sh1, n=4 for *SNHG29*-sh2). Right: CRISPR-CasRx using three sgRNAs (n=3, except n=2 for Rx3). Data normalized to day 0 and respective controls. (mean ± s.e.m; *P<0.05, **P<0.01, ***P<0.001; two-tailed, unpaired t-test) **C** *SNHG29* expression in M-07e cells via qRT-PCR. Left: CRISPRi knockdown (n=3 for sg1 and sg2, n=2 for sg3). Middle: RNAi knockdown (n=5 for sh1, n=4 for sh2). Right: CRISPR-CasRx knockdown (n=2, mean ± s.e.m.). Data normalized to respective controls. (mean ± s.e.m.; *P<0.05, **P<0.01, ***P<0.001; two-tailed, unpaired t-test) **D** Cell depletion at the final timepoint of fluorescence-based proliferation assay in M-07e cells using CRISPR-Cas9 targeting indicated SNORD loci and LRRC75A. Data normalized to day 0 and a non-targeting control sgRNA (n=3 biological replicates for LRRC75A, n=2 for other conditions, mean ± s.e.m.). **E** Cell depletion at the final timepoint of competitive proliferation assays targeting *SNHG29* in different AML cell lines. Left: CRISPRi-mediated depletion at day 18 (n=2 biological replicates, pooled data from 3 sgRNAs per cell line). Right: shRNA-mediated depletion at day 15 for M-07e cells and day 14 for other cell lines (n=4 in M-07e, n=2 for all other conditions, pooled data from 2 shRNAs per cell line). Data normalized to day 0 and controls (LUC-sg for CRISPRi, Ctrl-sh for RNAi). (mean ± s.e.m.; ***P<0.001.; two-tailed, unpaired t-test)

*SNHG29* hosts several C/D box snoRNAs (SNORD49A, SNORD49B, SNORD65). To test whether these embedded elements mediate the observed phenotype, we targeted conserved C/D motifs using CRISPR-Cas9 as previously described.^24^ In parallel, we noted that *SNHG29* knockdown modestly reduced expression of its neighboring coding gene, *LRRC75A* (**Figure S1A**). However, knockout of *LRRC75A* or the embedded snoRNAs did not affect proliferation, even accounting for variation in sgRNA efficiency (**Figure 2D**, **Figure S1B**). These findings indicate that the functional dependency resides in the *SNHG29* transcript itself, rather than its hosted snoRNAs or neighboring gene.

We next extended these findings to additional AML models. Only AMKL cell line M-07e exhibited growth inhibition upon *SNHG29* depletion (**Figure 2E**), whereas non-AMKL cell lines were unaffected, confirming the specificity of this dependency.

To test functional relevance in patient-derived material, we performed colony assays following *SNHG29* knockdown in primary AMKL blasts, which showed a significant reduction in colony-forming potential upon *SNHG29* depletion (**Figure 3A**). In competitive xenotransplantation assays, PDX cells transduced with *SNHG29* shRNA or a control (*LUC* shRNA) were co-injected into immunodeficient mice at a 1:1 ratio (**Figure 3B**). Endpoint analysis revealed that *SNHG29*-depleted cells were nearly eliminated (**Figure 3C**), demonstrating that *SNHG29* loss impairs leukemic propagation *in vivo*. These data provide an *in vivo* proof-of-concept for the therapeutic potential of targeting *SNHG29* in AMKL.

**Fig. 3.**
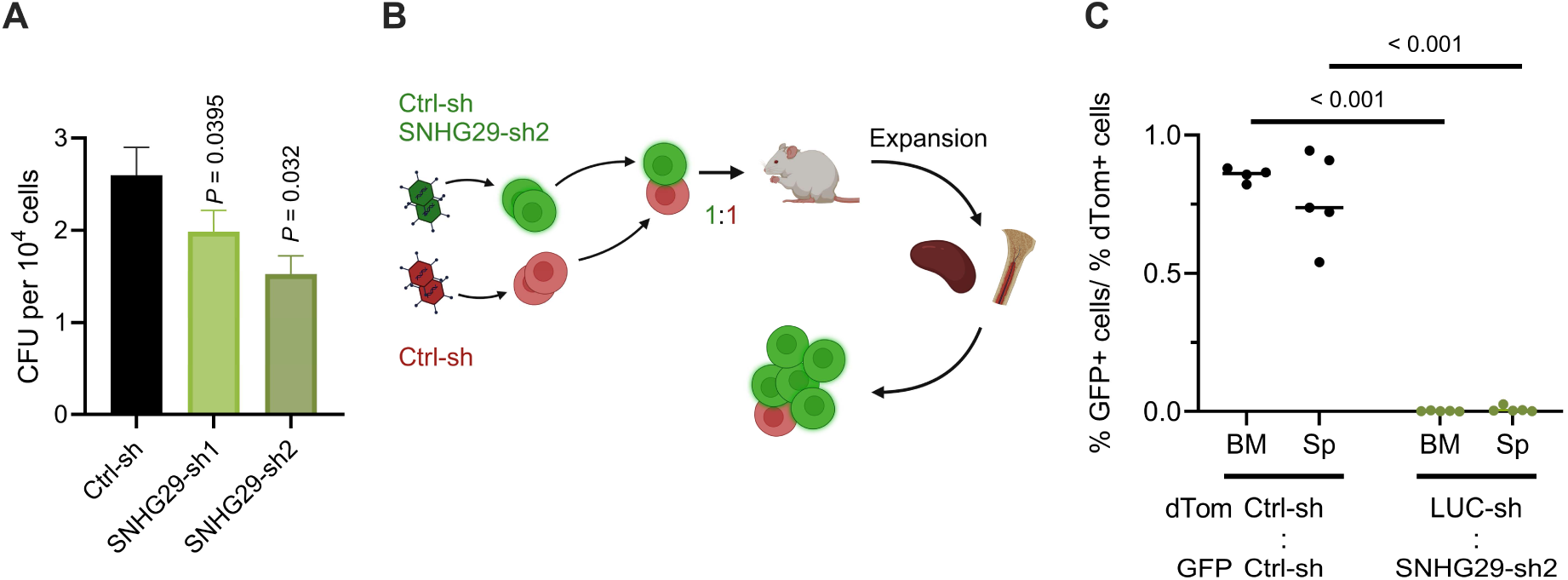
**A** Colony forming assay in AMKL PDX cells following *SNHG29* knockdown (n=3 biological replicates, mean ± s.e.m.; *P<0.05, **P<0.01) **B** Experimental setup for *in vivo* competition assays using AMKL PDX cells. **C** *In vivo* competition assay showing ratio of GFP to dTomato-labeled cells in bone marrow (BM) and spleen (Sp) of transplanted mice (n=5 biological replicates, mean ± s.e.m.; ***P<0.001, ****P<0.0001; two-tailed, unpaired t-test).

### SNHG29 interacts with the oncofetal RBP IGF2BP1

To investigate the mechanism by which *SNHG29* promotes AMKL proliferation, we performed RNA pulldown assays coupled with LC-MS/MS proteomic analysis (**Figure 4A**). Antisense oligos tiling the *SNHG29* locus were screened by RNase H accessibility to identify optimal capture sites (**Figure S1D**). Specific enrichment of *SNHG29* was confirmed by qRT-PCR in pulldown eluates (**Figure S1C**), and SDS-PAGE revealed distinct protein bands absent from controls (**Figure S1E**). LC-MS/MS identified 19 high-confidence proteins enriched in *SNHG29* pulldowns (**Figure 4C**, **Table S1**).

**Fig. 4.**
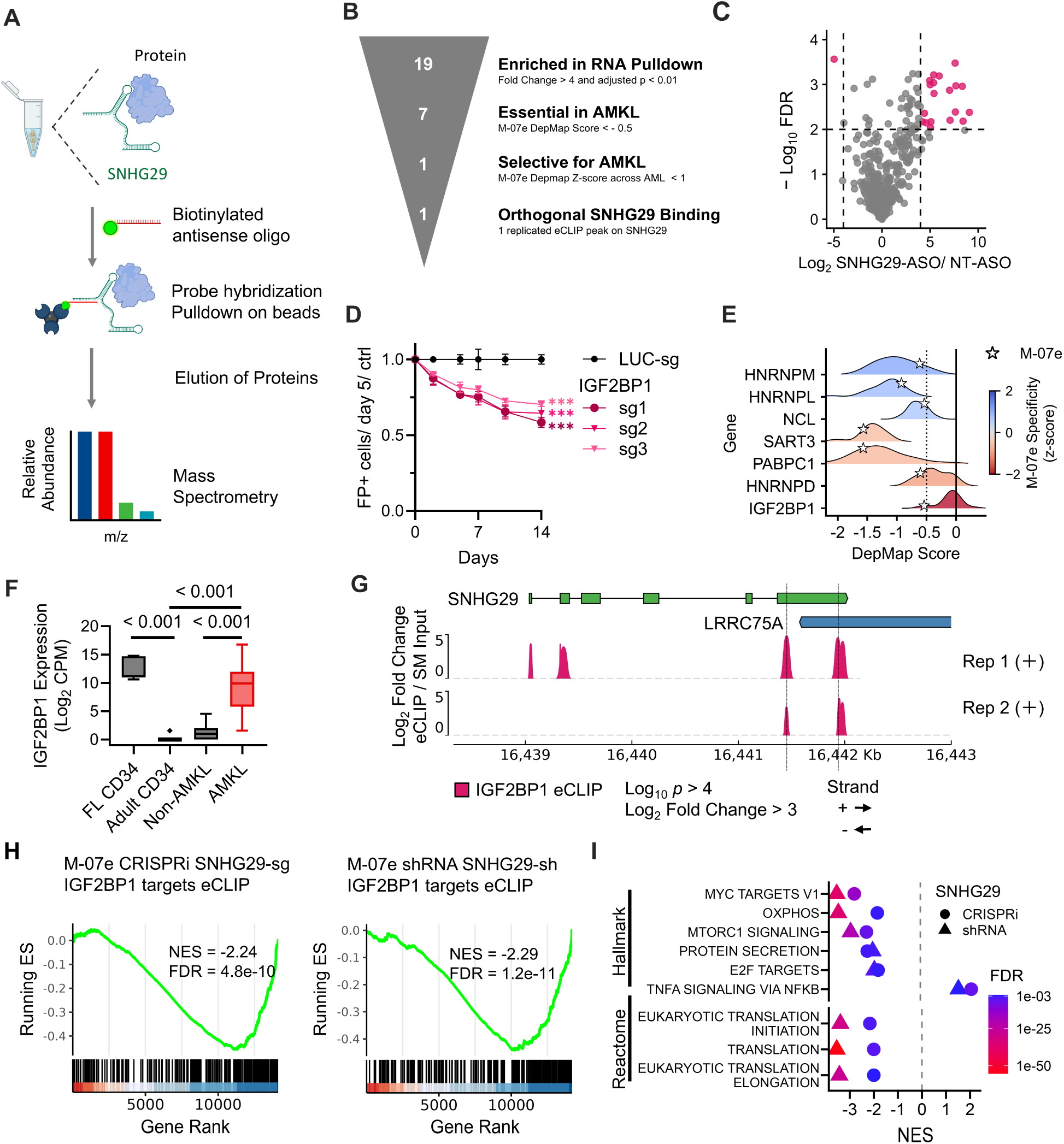
**A** Schematic representation of RNA pulldown strategy for identification of *SNHG29*-interacting proteins. **B** Schematic representation of our approach in determining a potentially mechanistic interaction with *SNHG29* in M-07e. **C** RNA pulldown of *SNHG29* in M-07e cells. Significantly enriched proteins (Log2 Fold Change > 4, adjusted P < 0.01) highlighted in pink (n=3 biological replicates, t-test, BH multiple testing correction, analysis by Perseus). **D** Fluorescence-based proliferation assays in M-07e cells with CRISPR-Cas9-mediated IGF2BP1 knockout using 3 sgRNAs. Data normalized to day 0 and respective controls. (mean ± s.e.m; *P<0.05, **P<0.01, ***P<0.001; two-tailed, unpaired t-test) **E** Dependency scores (DepMap 25Q3) of proteins enriched in *SNHG29* pulldown and essential in M-07e (DepMap score < −0.5) across 29 AML cell lines. Specificity Z-scores were calculated across AML cell lines. **F** Expression of IGF2BP1 in RNA sequencing datasets from fetal liver CD34+ cells (FL CD34, n=5), peripheral blood mobilized CD34+ cells (PB CD34, n=8), AMKL (n=19) and non-megakaryoblastic AML (non-AMKL) (n=130) patient samples. Data shown are Log2 transformed TPM. Box plots show medians, boxes and whiskers according to the Tukey method. ***FDR<0.001 (LIMMA-voom). **G** eCLIP peak data showing IGF2BP1 binding at the *SNHG29* locus in K562. Plus strand peaks with - Log10 P > 4 and Log2 fold change > 3 over size-matched input are displayed. No significant peaks were detected on the minus strand. (n=2) **H** Gene Set Enrichment Analysis (GSEA) of the top 200 IGF2BP1 eCLIP targets following *SNHG29* knockdown in M-07e cells using CRISPRi with two different sgRNAs (left) or two different shRNAs (right) analyzed compared to the respective non-targeting control. Genes from RNA-seq experiments (n=3) were ranked by t-statistic. NES, Normalized Enrichment Score; FDR, False Discovery Rate; ES, Enrichment Score. **I** GSEA of Hallmark and Reactome gene sets following *SNHG29* knockdown via shRNA and CRISPRi (n=3). Gene sets with the highest averaged NES between shRNA and CRISPRi conditions are shown.

To prioritize functionally relevant interactors, we integrated proteomic enrichment, DepMap^25^ CRISPR dependency data, and eCLIP binding profiles (**Figure 4B**). Of the 19 candidates (fold change >4, FDR <0.01), seven were essential for M-07e survival (DepMap score <-0.5, **Figure S2A**). Of these, only IGF2BP1 showed selective dependency in M-07e (DepMap Z-score <-1, **Figure 4E**). Most other proteins represented common essential splicing factors. Cross-referencing with eCLIP datasets^26^ showed IGF2BP1 binding on *SNHG29* exons (**Figure 4G**). This integrative filtering approach thus pinpointed IGF2BP1 as the principal mechanistic partner of *SNHG29* in AMKL.

### The SNHG29–IGF2BP1 interaction drives AMKL dependency

CRISPR knockout of IGF2BP1 in M-07e cells phenocopied *SNHG29* loss, causing robust growth inhibition (**Figure 4D**). In DepMap^25^ data, this dependency was restricted to AMKL-derived lines (M-07e, CMK, and the erythroid-like OCI-M2), whereas other AML subtypes were unaffected (**Figure 4E**). eCLIP analysis^26^ confirmed two IGF2BP1 binding sites on the terminal exon of *SNHG29* (**Figure 4G**). Notably, *IGF2BP1* is expressed in fetal hematopoietic progenitors, silenced in adult hematopoiesis, and retained specifically in AMKL (**Figure 4F**). More broadly, AMKL blasts exhibited upregulation of fetal hematopoietic gene signatures compared to non-AMKL AML subtypes (**Figure S1H**).

RNA-seq data revealed strong correlation between *SNHG29* expression and IGF2BP1 target genes, including *MYC* and *E2F1*, across AMKL samples (**Figure S2B**), whereas *IGF2BP1* transcript levels alone did not correlate with *SNHG29*. These findings suggest that *SNHG29* modulates *IGF2BP1’s* functional output rather than its abundance.

To assess the transcriptional consequences of *SNHG29* loss, we performed RNA-seq following CRISPRi or shRNA-mediated depletion of *SNHG29*. Expression of IGF2BP1 target mRNAs (defined by eCLIP) was consistently downregulated upon *SNHG29* knockdown (**Figure 4H**), as were of *MYC* and *E2F* target pathways and translational control programs (**Figure 4I**). Collectively, these data suggest that *SNHG29* supports AMKL growth by stabilizing IGF2BP1-bound transcripts and maintaining downstream pathways.

### Pharmacologic inhibition of IGF2BP1 selectively impairs AMKL viability

Given the dependency of AMKL cells on the *SNHG29*–*IGF2BP1* axis, we hypothesized that direct pharmacologic inhibition of IGF2BP1 would recapitulate the effects of *SNHG29* depletion. Treatment of AMKL cells with BTYNB, a small molecule that blocks IGF2BP1RNA binding, resulted in potent and selective cytotoxicity in M-07e cells, with an LC_50_ of 1.9 µM (**Figure 5A, B**). Non-AMKL AML cell lines displayed markedly reduced sensitivity, indicating subtype-specific vulnerability.

**Fig. 5.**
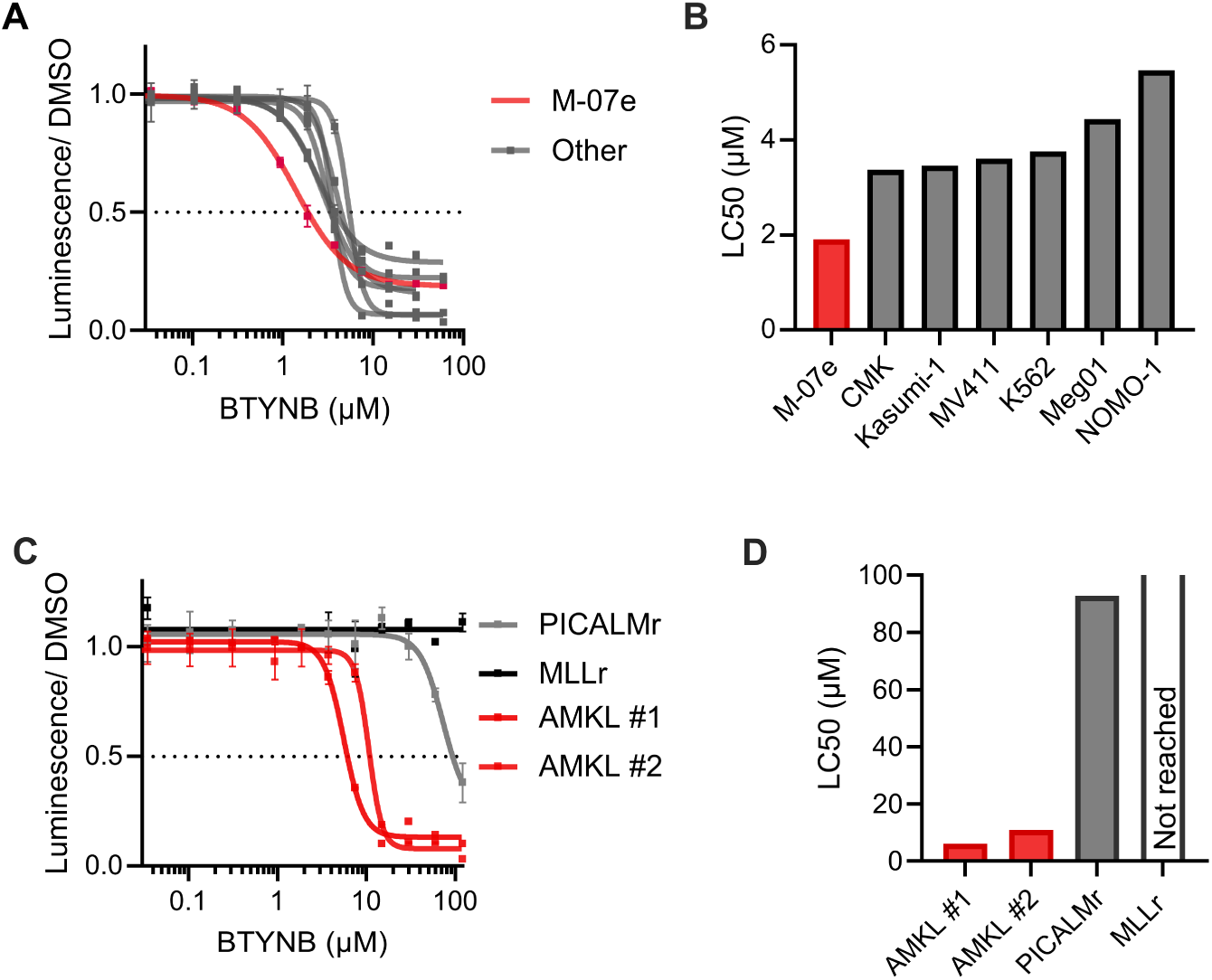
**A** Dose-response curves for the AMKL cell line M-07e and non-AMKL cell lines after 72-hour treatment with the IGF2BP1 inhibitor BTYNB. Viability was measured using a luminescence-based cell viability assay and normalized to vehicle-treated controls (n=3 biological replicates, mean ± s.e.m.). **B** LC_50_ values for BTYNB in the indicated cell lines, derived from non-linear regression (n=3 biological replicates). **C** Dose-response curves for AMKL and non-AMKL patient-derived blasts after a 5-day treatment with BTYNB. Viability was measured and normalized as in (A) (n=3 biological replicates, mean ± s.e.m.). **D** Absolute LC_50_ values for BTYNB in AMKL and non-AMKL patient-derived blasts, determined by non-linear regression (n=3 biological replicates).

We next tested BTYNB in patient-derived AMKL blasts, which showed pronounced and dose-dependent cytotoxicity (**Figure 5C**). In contrast, non-AMKL pediatric AML samples, including a *KMT2A*-rearranged case, were resistant to treatment (**Figure 5D**). These results demonstrate that AMKL cells are selectively sensitive to pharmacologic disruption of IGF2BP1, providing a proof-of-concept for targeting the *SNHG29*–IGF2BP1 axis as a therapeutic strategy in this disease.

Together, these results identify *SNHG29* as a lineage-specific dependency that sustains AMKL proliferation through its interaction with the oncofetal RBP IGF2BP1. *IGF2BP1* transcript levels did not correlate with *SNHG29* expression, yet IGF2BP1 target mRNAs were consistently downregulated upon *SNHG29* depletion, suggesting that *SNHG29* enhances IGF2BP1-mediated mRNA stabilization rather than regulating IGF2BP1 abundance. The selective activation of this developmental RNA-RBP circuit in AMKL creates a lineage-restricted vulnerability that can be therapeutically targeted. Pharmacologic inhibition of IGF2BP1 with BTYNB recapitulates the effects of *SNHG29* depletion, demonstrating the translational feasibility of disrupting lncRNA–RBP interactions in this aggressive leukemia.

These results define a developmental RNA-RBP dependency unique to AMKL and establish a mechanistic framework for understanding how oncofetal networks sustain malignant growth – setting the stage for therapeutic exploration of the *SNHG29*–IGF2BP1 axis.

## Discussion

This study identifies *SNHG29* as a previously unrecognized and essential lncRNA in AMKL, revealing a lineage-specific dependency that defines a new therapeutic vulnerability. Using an integrated strategy that combined CRISPRi screening, multiple knockdown approaches, and functional validation in vitro and in vivo, we demonstrate that *SNHG29* is indispensable for leukemic proliferation and survival. These results position *SNHG29* as a crucial regulator of AMKL maintenance and a potential therapeutic target in this high-risk leukemia with limited treatment options.

*SNHG29* depletion caused profound loss of cell viability, reduced clonogenic growth, and conferred a competitive disadvantage in PDX models, underscoring its functional importance. The specificity of this dependency for AMKL, and its absence in other AML subtypes, highlights a lineage-restricted vulnerability shaped by developmental context. Such selectivity suggests that targeting *SNHG29* or its effector pathways could impair leukemic growth without broadly affecting normal hematopoietic cells.

Mechanistically, *SNHG29* functions through direct interaction with the oncofetal RBP IGF2BP1, a post-transcriptional regulator of mRNAs encoding oncogenic drivers such as MYC and E2F1^27–30^. Although IGF2BP1 has established roles in tumorigenesis, its contribution to pediatric AMKL has not been previously defined. Our findings reveal that *SNHG29* is required to sustain IGF2BP1 activity and stabilize its RNA targets. Loss of *SNHG29* phenocopies IGF2BP1 inhibition, leading to downregulation of IGF2BP1-bound transcripts and suppression of downstream proliferative pathways. These results uncover a previously unappreciated post-transcriptional circuit in which a lncRNA cooperates with an oncofetal RBP to maintain leukemic growth.

*SNHG29* enhances IGF2BP1 function rather than antagonizing it, consistent with prior observations that lncRNAs can act as positive regulators of IGF2BP family proteins. Similar mechanisms have been described for THOR and KB-1980E6.3, which promote IGF2BP1-mediated mRNA stabilization in solid tumors.^31–33^ This interaction is specific to AMKL, representing a distinct oncofetal RNA axis reactivated in hematologic malignancy. This context-dependent interaction underscores the modularity of lncRNA function and its capacity to reinforce oncogenic pathways through protein partners selectively expressed in developmental lineages.

Our data also provide a mechanistic framework to reconcile previous reports describing *SNHG29* as a tumor suppressor in *FLT3-ITD*–positive AML, where it interacts with EP300.^34^ We propose that *SNHG29*’s function is dictated by the cellular context and availability of lineage-specific cofactors. In the oncofetal environment of AMKL, characterized by high IGF2BP1 expression, *SNHG29* is co-opted into an oncogenic role. This plasticity exemplifies how lncRNAs can adopt distinct, even opposing, functions depending on transcriptional and epigenetic context.

The identification of a selective *SNHG29*–IGF2BP1 dependency offers immediate translational potential. Direct targeting of lncRNAs remains technically challenging, but their protein effectors are more amenable to pharmacologic inhibition. We show that AMKL cells are selectively sensitive to BTYNB, a small-molecule inhibitor that disrupts IGF2BP1–RNA interactions.^35^ Treatment with BTYNB potently reduced viability in AMKL cell lines and patient-derived blasts while having limited effects on other AML subtypes. This selective sensitivity supports a model in which AMKL cells are addicted to the *SNHG29*–IGF2BP1 axis and demonstrates that pharmacologic interference with RNA–protein complexes can be therapeutically viable.

Our findings also contribute to the evolving understanding of lncRNA-mediated oncogenesis. Several pro-leukemic lncRNAs regulate gene expression at the transcriptional level,^36^ exemplified by HOTTIP,^37^ or modulate cytoplasmic signaling pathways such as HOXA10-AS.^38^ *SNHG29* belongs to a distinct class of lncRNAs that act at the post-transcriptional level, directly modulating RBP activity to control mRNA fate. This mechanism highlights the functional versatility of lncRNAs as integrators of RNA metabolism and cancer signaling. Moreover, the activation of an oncofetal RBP in AMKL suggests that malignant cells can repurpose fetal regulatory networks to sustain proliferation, a concept increasingly recognized across pediatric cancers.

The role of IGF2BP1 in AMKL aligns with its role during fetal hematopoiesis and indicates that developmental gene expression programs are aberrantly retained in this leukemia. By partnering with *SNHG29*, IGF2BP1 reinstates a fetal post-transcriptional landscape that promotes leukemic maintenance. This model provides a developmental rationale for AMKL pathogenesis and reveals how fetal regulators can be activated to sustain malignancy. Understanding how these programs are re-engaged in cancer will be key to defining selective therapeutic opportunities.

The context specificity of the *SNHG29*–IGF2BP1 dependency raises further questions about its regulation. Both factors are upregulated in AMKL, potentially driven by developmental transcription factors or epigenetic reprogramming unique to AMKL. Identifying these upstream regulators could explain how this oncofetal circuit is selectively activated in the megakaryoblastic lineage and may reveal additional therapeutic entry points. It will also be important to determine whether similar lncRNA–RBP partnerships operate in other pediatric or developmental cancers.

The selective vulnerability of AMKL cells to IGF2BP1 inhibition suggests that such lineage-restricted dependencies could define synthetic lethal opportunities in cancer. The developmental confinement of this axis may also reduce toxicity risks, as adult tissues largely silence both *SNHG29* and IGF2BP1. These features make the *SNHG29*–IGF2BP1 pathway particularly attractive for targeted therapy, offering a rare combination of mechanistic specificity and potential therapeutic index.

While this study establishes *SNHG29* as a key regulator of IGF2BP1 activity in AMKL, several questions remain. Structural elucidation of the *SNHG29*–IGF2BP1 interface will be important to define the molecular basis of their cooperation and could inform rational drug design. Expanded analyses across primary patient samples are needed to validate the generality of this dependency and its potential relevance in other hematologic malignancies. Finally, BTYNB establishes proof-of-concept for pharmacologic disruption of IGF2BP1–RNA interactions but will require substantial optimization and comprehensive pharmacodynamic and safety evaluation to enable clinical translation.

In summary, our work uncovers a developmental RNA circuit in which the lncRNA *SNHG29* cooperates with the oncofetal RBP IGF2BP1 to sustain leukemic gene expression programs in AMKL. This axis defines a lineage-restricted dependency that can be therapeutically exploited. By elucidating a post-transcriptional mechanism linking fetal RNA regulation to leukemic maintenance, our findings expand the paradigm of lncRNA–RBP cooperation in cancer and highlight the potential of targeting RNA-based vulnerabilities in pediatric malignancies.

## Acknowledgements

This work was supported by the following funding: J.H.K. – European Research Council Horizon 2020 program (#714226), *Hilfe für krebskranke Kinder Frankfurt e.V.* (C^3^OMBAT-AML) and the German Research Foundation (DFG; FOR 5433 *RNA in Focus*; KL-2371/7-1468534282); S.H. – DFG (FOR 5433; 468534282). DH is supported by the Frankfurt Foundation for Children with Cancer.

Illustrations were created with BioRender.com.

## Author contributions

R.W. and R.B. performed experiments and analyzed the results. R.W. and M.N. conducted bioinformatic analyses. D.H. and J.H.K. designed and supervised the study. R.W., R.B., and J.H.K. crafted the manuscript. All authors revised and approved the manuscript.

## Methods

### Lentiviral Vectors

To perform targeted genetic perturbations, we utilized several lentiviral systems. Stable cell lines expressing the necessary effector proteins were generated with pLKO5d.SFFV.dCas9-KRAB.MeCP2.P2A.BSD for CRISPRi, pLKO5d.EFS.SpCas9.P2A.BSD (Addgene #57821) for knockout, or pLKO5d.SFFV.hfCas13d.p2a.BSD for RNA targeting. Individual sgRNAs, designed with the CCTop^39^ platform, were delivered via the SGL40C.EFS.E2Crimson vector (Addgene #100894). For RNA interference, we designed shRNA sequences using the miR-N tool^40^ and cloned them into either the SIN40C.SFFV.eGFP.miR30n (Addgene #169278) or SIN40C.SFFV.dTomato.miR30n (Addgene #169277) backbone via BsmBI digestion. The sgRNA library for the CRISPRi screen was expressed from the SGL40C.EFS.dTomato vector (Addgene #89395). To perturb IGF2BP1, previously validated sgRNAs graciously provided by Prof. Hüttelmaier were utilized. For LRRC75A knockout, sgRNA sequences were derived from the GeCKO v2 library.^41^ In all perturbation experiments, constructs targeting firefly luciferase were used as non-targeting negative controls.

### Lentivirus Production

Lentiviral particles were generated via PEI-mediated co-transfection of HEK293T cells with the respective expression vector alongside packaging plasmids pMD2.G (Addgene #12259) and psPAX2 (Addgene #12260). Virus-containing supernatant was harvested and concentrated by ultracentrifugation. Target cells were then transduced by incubation with the concentrated virus in media supplemented with 5 µg/ml Polybrene (Sigma-Aldrich).

### CRISPR Library Design, Cloning and Screening

We employed a custom CRISPRi library designed to target a comprehensive list of lncRNAs assembled from GENCODE v25(03/2016 release),^42^ LNCipedia 4.0 (05/2016 release)^43^ and NONCODE v4 (01/2014 release),^44^ that was previously developed by our group.^45^ The library comprises three to nine sgRNAs per target, tiling the 250 bp window immediately downstream of the annotated transcription start site. The final library incorporated non-targeting controls against luciferase and positive depletion controls targeting essential genes such as *MYC*, *MYB*, and *RPL9*.

Library construction involved cloning a pooled set of synthesized oligonucleotide spacers (Integrated DNA Technologies) into the SGL40C.EFS.dTomato vector (Addgene 89395) via BsmBI restriction sites. After transformation into XL1-Blue supercompetent cells (Agilent 200236) and plating, library representation was confirmed by colony counting. The screen itself was conducted in dCas9-KRAB.MeCP2-expressing cell lines. Cells were transduced with the lentiviral library at 30% efficacy and then cultured for 18 population doublings, ensuring a library representation of at least 1000-fold was maintained throughout. To prepare the library for sequencing, genomic DNA was extracted from samples at the beginning and end of the screen (JetFlex™ Genomic DNA Purification Kit, Thermo Fisher Scientific). The integrated sgRNA cassettes were then amplified by PCR (NEBNext® High-Fidelity 2x PCR Master Mix, New England Biolabs), gel-purified (GeneJET Gel Extraction Kit, Thermo Fisher Scientific), and sequenced on an Illumina HiSeq 2000 instrument. To identify AMKL-specific lncRNA dependencies, we analyzed the resulting 50 bp single-end reads from all cell lines using the MAGeCK MLE algorithm.^46^

### Fluorescence-based Proliferation Assay

The impact of individual gene perturbations on cell proliferation was assessed using competitive proliferation assays. Stable cell lines expressing the relevant Cas effector or wild-type cells (for RNAi) were transduced to create a mixed population of perturbed (fluorescent) and unperturbed cells. The proportion of fluorescent cells in the culture was then monitored by flow cytometry every 2-3 days for up to three weeks. The growth phenotype was quantified by normalizing the percentage of fluorescent cells at each time point against its starting value on day 0 and against a non-targeting (LUC) control.

### Quantitative Real-Time PCR

Total RNA was extracted from cells between days 3 and 7 post-transduction using Quick-RNA^TM^ Microprep or Miniprep Kits (Zymo Research) and contaminating genomic DNA was removed using on-column DNase digestion (Zymo Research) or the TURBO DNA-free^TM^ Kit (Thermo Fisher Scientific). cDNA was synthesized using the High-Capacity cDNA Reverse Transcription Kit (Applied Biosystems). qPCR was then performed using SYBR™ Select Master Mix on a Quantstudio 5 or StepOnePlus^TM^ Real-Time PCR cycler, and expression levels were normalized to the housekeeping gene B2M. For cellular fractionation experiments the procedure was performed as previously described,^47^ with B2M and GAPDH serving as cytoplasmic controls, and XIST and TERC as nuclear controls.

### Flow Cytometry and Cell Sorting

Flow cytometry data were acquired on a CytoFLEX B4-R3-V5 or CytoFLEX S V4-B2-Y4-R3 instrument (Beckman Coulter) and analyzed using FlowJo™ v10.9 software (BD Biosciences). For cell sorting, a FACSAria™ II instrument running FACSDiva™ software (BD Biosciences) was utilized.

### In Vivo Xenotransplantation Assay

To assess the *in vivo* effects of *SNHG29* knockdown, we employed a competitive xenotransplantation assay using murine xenograft models of AML according to previously established protocols.^38,48^ *In vivo*-expanded PDXs were transduced with either GFP or dTomato shRNA vectors. Two experimental conditions were established to control for effects of fluorophore expression, GFP LUC-sh versus dTomato LUC-sh and GFP LUC-sh versus dTomato *SNHG29*-sh2. The two populations were mixed at a 1:1 ratio, and one to two million total cells were administered via tail vein into 8-10 week old MISTRG mice that had received 2.5 Gy irradiation. We monitored leukemic engraftment by analyzing peripheral blood every two weeks via flow cytometry. At humane endpoints, mice were euthanized, and cells were harvested from bone marrow and spleen for final analysis. The competitive disadvantage caused by *SNHG29* knockdown was determined by comparing the final ratio of the dTomato and GFP populations. All animal experiments were performed at the Goethe University Frankfurt animal facility under protocols approved by the Regierungspräsidium Darmstadt.

### RNase H Accessibility Assay

To identify accessible regions on the *SNHG29* transcript, we designed a series of 21 nt antisense DNA oligonucleotides (ASOs) tiling its sequence, excluding those with likely off-target binding as predicted by BLAST. The assay was performed following a modified protocol from Neumann et. al.^49^ Briefly, lysates from M-07e cells were adjusted to a final buffer composition of 60 mM NaCl, 50 mM Tris-HCl pH 8, 75 mM KCl, 3 mM MgCl₂, and 10 mM DTT. The lysates were incubated with 100 pmol of each ASO for 2 h at 4°C before the addition of RNase H. Following a 20-minute incubation at 37°C to induce cleavage, total RNA was isolated for qRT-PCR analysis using primers flanking each ASO target site.

### RNA Affinity Purification

RNA affinity purification was performed based on a protocol previously described by Neumann et. al.^49^ For each experimental condition, 20 million M-07e cells were washed and lysed in lysis buffer (Tris-HCl 50mM, NaCl 150mM, EDTA 1mM, 1% NP-40, Roche c0mplete protease inhibitor). The resulting lysate was pre-cleared with streptavidin beads for 2 h at 4°C, then incubated with 200 pmol of 2’O-Me-RNA antisense oligonucleotides for 1 h at 37°C. To capture the RNA-protein complexes, we used streptavidin beads that had been pre-blocked with 0.2 mg/ml each of ytRNA (Thermo Fisher Scientific) and glycogen (Thermo Fisher Scientific).

The beads were subjected to five washes in wash buffer (50 mM Tris-HCl pH8, 50 mM NaCl, 0.05% NP-40, Roche c0mplete Protease inhibitor), excluding protease inhibitors and NP-40 during the final wash step. Elution was performed by incubating the beads for 30 minutes at 37°C in 50 µL of elution buffer (10 mM D-Biotin, 10 mM Tris-HCl, 50 mM NaCl). Eluates were analyzed by qRT-PCR and relative *SNHG29* enrichment calculated over input samples. SDS-PAGE was stained using the SilverQuest silver staining kit (Thermo Fisher Scientific).

Eluates were trypsinated and LC-MS/MS was performed on a Q Exactive Plus instrument (Thermo Fisher Scientific) equipped with an ultra-high performance liquid chromatography unit (Thermo Scientific Dionex Ultimate 3000). For data analysis using t-test and multiple testing correction MaxQuant 2.0.3.0^50^, Perseus 2.0.7.0^51^ and Excel (Microsoft Office 2016) were used. The UniProt^52^ human reference proteome database (October 2022, 102557 entries) was used to identify peptides and proteins. Reverse identifications and common contaminants were removed. Missing values were replaced by background values from normal distribution. Proteins with Log_2_FoldChange > 4 and adjusted P < 0.01 in the *SNHG29*-ASO vs NT-ASO condition were considered for downstream analysis.

### DepMap Dependency Analysis

CRISPR dependency scores for *SNHG29* pulldown enriched proteins were retrieved from the DepMap portal (25Q3 release). Analysis included all 29 AML cell lines available in DepMap (**Table S6**). Genes were classified as essential in M-07e if their dependency score was < −0.5. To identify selective dependencies in M-07e, Z-scores were calculated by standardizing each gene’s dependency score in M-07e against the mean and standard deviation across all 29 AML cell lines. Genes with a Z-score < −1 were considered selectively essential in M-07e.

### Enhanced UV Crosslinking and Immunoprecipitation (eCLIP) Analysis

eCLIP data for proteins showing significant binding in the RNA affinity purification was retrieved from the Encode consortium. Data consisted of 2 replicates in HepG2, K562 or both. K562 experiments were preferred where available. Preprocessing steps, including alignment to hg38 and peak calling using CLIPper, were previously carried out by the ENCODE consortium. Individual replicate peaks were mapped to Ensembl version 108^53^ transcripts using bedtools.^54^ To orthogonally validate protein binding to *SNHG29*, specific criteria were applied: a peak had to be present on a *SNHG29* exon within 100bp in both replicates, with a Log_2_ Fold Change > 3 over size-matched input and a -Log_10_ adjusted P value > 4. PyGenomeTracks was used to visualize eCLIP peaks on the *SNHG29* transcript. Target genes were classified as significantly bound if a reproducible peak was found in the IDR peak set. Both intronic and exonic binding was considered. Bound genes were ranked by Log_2_ Fold Change. Gene sets were created after sorting by Log_2_ Fold Change to select 250 genes for gene set enrichment analysis (GSEA).

### RNA Sequencing

For transcriptomic analysis, total RNA was extracted from cells on days 3 (shRNA) or 7 (CRISPRi) post-transduction using the RNeasy Plus Mini Kit (Qiagen), with timepoints selected based on depletion dynamics. Novogene Company, Ltd. conducted rRNA depletion and sequencing on an Illumina NovaSeq 6000 (150 bp paired-end). The raw sequencing data were processed using the nf-core/rnaseq pipeline^55,56^ with default settings. Briefly, reads were aligned to the hg38 reference genome using STAR^57^ and quantified using Salmon^58^ against the Ensembl transcriptome version 108.^53^ Existing batch effect from flow cell lane was corrected using Combat-seq,^59^ setting the experimental condition as the variable of interest. Differential expression analysis was performed separately for CRISPRi and shRNA in R using DESeq2.^60^ To account for the large difference in knockdown efficacy between *SNHG29*-sh1 and -sh2 (Figure 2D), Log_2_ *SNHG29* relative expression as determined by qRT-PCR was incorporated in the linear model. Log fold changes were shrunk using apeGLM,^61^ according to best practice. Genes were ranked by t-statistic for GSEA. Gene sets Hallmark and Reactome were retrieved from MSigDB v2024.1.Hs.^62^ Gene sets of RBP targets were derived from eCLIP analysis as described above. Gene set enrichment analysis was carried out using fgsea in R.^63^

Patient RNA-Sequencing data was aligned and quantified as above. Genes with less than 10 counts in more than two thirds of samples of all molecular subtypes were filtered out. Length-corrected counts were normalized and variance stabilized using LIMMA-voom.^64^ Pearson correlation to *SNHG29* expression was calculated on Log_2_ transformed counts per million and multiple testing corrected using the Benjamini-Hochberg procedure. Differential expression analysis was performed using LIMMA. To assess fetal hematopoietic gene expression, two complementary gene sets were used. A human fetal signature was derived from differentially expressed genes between human fetal liver and adult HSPCs, and a mouse fetal signature was derived from mouse fetal liver versus adult HSPCs with gene symbols converted to human orthologs. Signature enrichment scores were calculated using GSVA.^65^

### In Vitro Dose-Response Assay

To assess the efficacy of BTYNB, leukemia cell lines and PDX cells were seeded in 96-well plates. The cells were treated with serial dilutions of BTYNB or a DMSO vehicle control. Cell lines were incubated for 3 days, while PDX cells were cultured for 5 days. Cell viability was subsequently measured using the CellTiter-Glo® Luminescent Cell Viability Assay (Promega) following the manufacturer’s instructions. Luminescence was recorded on a GloMax® Multi+ detection system (Promega). Viability data were normalized to vehicle-treated control wells. LC_50_ values, defined as the drug concentration required to reduce cell viability to 50%, were then calculated from dose-response curves using nonlinear regression analysis in GraphPad Prism 9.

### CRISPR-Cas9 Indel Validation

The efficiency of CRISPR-Cas9-mediated gene editing was confirmed by analyzing insertions and deletions (indels). Genomic DNA was isolated from cells 3 days post-transduction using the Quick-DNATM Miniprep Kit (Zymo Research). A PCR amplicon of approximately 500 bp flanking the sgRNA target site was generated and subjected to Sanger sequencing. The resulting sequence trace was compared to that from wild-type cells using the TIDE online tool^66^ to quantify the percentage of edited alleles.

### Quantifications and Statistical Analysis

Data from the CRISPRi screen were analyzed using the MAGECK^46^ algorithm. For all other experiments, statistical analyses were performed in GraphPad Prism 9. Two-tailed, unpaired t-tests were used for comparisons between groups, except where a ratio paired t-test was employed (Figure S1B). RNA affinity purification data were analyzed by student’s t-test and multiple testing correction by Benjamini-Hochberg procedure in MaxQuant 2.0.3.0,^50^ Perseus 2.0.7.0^51^ and Excel (Microsoft Office 2016). Differential expression analysis of RNA-Seq was performed using DESeq2^60^ and LIMMA,^64^ for *SNHG29* knockdown and AML patient data, respectively. The correlation between groups was analyzed by Pearson correlation and differences between correlation coefficients were compared by a two-tailed t-test after Fisher z-transformation. All data are presented as mean ± s.e.m., with sample sizes indicated in the figure legends. A P-value of <0.05 was considered statistically significant. Sample sizes were not predetermined by statistical methods.

## Experimental Model and Study Participant Details

### Cells and Cell Culture

Patient-derived AML cells, previously expanded *in vivo*, were thawed and cultured for 24–48 hours prior to experiments in StemSpan SFEM (STEMCELL Technologies) supplemented with 1% penicillin/streptomycin, a cytokine cocktail (50 ng/ml SCF, 50 ng/ml FLT3L, 10 ng/ml IL6, 2.5 ng/ml IL3, 10 ng/ml TPO; Peprotech), and small molecules (750 nM SR1, 35 nM UM171; STEMCELL Technologies). At 72 hours post-transduction, cells were either harvested and sorted for subsequent xenotransplantation and colony-forming assays or continuously cultured for *in vitro* competition assays. Human myeloid leukemia cell lines K562, THP-1, ML-2, M-07e, KASUMI-1, NOMO-1 and SKNO-1 were procured from DSMZ (Braunschweig, Germany). All cell lines were maintained as per the supplier’s recommendations and were subjected to routine testing for Mycoplasma contamination. Transductions were performed with the addition of 5 µg/ml Polybrene (Sigma-Aldrich).

### Animal Studies

Animal experiments utilized M-CSFh/h IL-3/GM-CSFh/h SIRPah/h TPOh/h RAG2-/-IL2Rg-/-(MISTRG) immunodeficient mice (Regeneron Pharmaceuticals), which were maintained in a pathogen-free facility in individually ventilated cages with access to autoclaved food and water in a 12-hour light/dark cycle. All animal procedures were approved by the Regierungspräsidium Darmstadt and conducted at Goethe University Frankfurt.

### Human Participants

All human samples were collected with informed consent from participants or their legal guardians, under protocols approved by the ethics committee of the Martin Luther University Halle-Wittenberg. Primary acute myeloid leukemia (AML) samples were sourced from Goethe University Frankfurt, the AML-BFM Study Group (Essen, Germany), and the Department of Hematology, Hemostasis, Oncology and Stem Cell Transplantation at Hannover Medical School.

## Resource Availability

### Data and Code Availability

Raw CRISPR screening and RNA-seq data will be made available upon journal publication in the Gene Expression Omnibus (GEO). Data on CRISPR dependencies in AML cell lines were sourced from the DepMap Project.^25^ eCLIP datasets ENCSR125CLF and ENCSR975KIR were retrieved from ENCODE.^67^ Raw RNA-Seq data of AML patients is not published with this paper. The previously published^6^ microarray data of normal hematopoietic cells and pediatric AML used in this study can be accessed at https://ag-klusmann.shinyapps.io/lncScape/.

This study relied on standard computational algorithms detailed in the Methods section. Any original code or further details necessary for reanalysis will be made available by the author upon reasonable request.

## Supplemental tables

Table S1: CRISPRi lncRNA library.

Table S2: Clinical and genetic characteristics of patient cells used in this study.

Table S3: sgRNA and shRNA spacer sequences.

Table S4: PCR primers.

Table S5: Antisense oligonucleotides.

## References

1. Mercer, T. R., Dinger, M. E. & Mattick, J. S. Long non-coding RNAs: insights into functions. Nat. Rev. Genet. 10, 155–159 (2009).

2. Ponting, C. P., Oliver, P. L. & Reik, W. Evolution and Functions of Long Noncoding RNAs. Cell 136, 629–641 (2009).

3. Cabili, M. N. et al. Integrative annotation of human large intergenic noncoding RNAs reveals global properties and specific subclasses. Genes Dev. 25, 1915–1927 (2011).

4. Hon, C.-C. et al. An atlas of human long non-coding RNAs with accurate 5′ ends. Nature 543, 199–204 (2017).

5. Tsai, M.-C. et al. Long Noncoding RNA as Modular Scaffold of Histone Modification Complexes. Science 329, 689–693 (2010).

6. Zhao, J., Sun, B. K., Erwin, J. A., Song, J.-J. & Lee, J. T. Polycomb Proteins Targeted by a Short Repeat RNA to the Mouse X Chromosome. Science 322, 750–756 (2008).

7. Kretz, M. et al. Control of somatic tissue differentiation by the long non-coding RNA TINCR. Nature 493, 231–235 (2013).

8. Mattick, J. S. et al. Long non-coding RNAs: definitions, functions, challenges and recommendations. Nat. Rev. Mol. Cell Biol. 24, 430–447 (2023).

9. Kopp, F. & Mendell, J. T. Functional Classification and Experimental Dissection of Long Noncoding RNAs. Cell 172, 393–407 (2018).

10. Luo, M. et al. Long non-coding RNAs control hematopoietic stem cell function. Cell Stem Cell 16, 426–438 (2015).

11. Paralkar, V. R. et al. Lineage and species-specific long noncoding RNAs during erythromegakaryocytic development. Blood 123, 1927–1937 (2014).

12. Ng, M., Heckl, D. & Klusmann, J.-H. The Regulatory Roles of Long Noncoding RNAs in Acute Myeloid Leukemia. Front. Oncol. 9, (2019).

13. Papaioannou, D. et al. The long non-coding RNA HOXB-AS3 regulates ribosomal RNA transcription in NPM1-mutated acute myeloid leukemia. Nat. Commun. 10, 5351 (2019).

14. Bill, M. et al. Expression and functional relevance of long non-coding RNAs in acute myeloid leukemia stem cells. Leukemia 33, 2169–2182 (2019).

15. de Rooij, J. D. E. et al. Pediatric non-Down syndrome acute megakaryoblastic leukemia is characterized by distinct genomic subsets with varying outcomes. Nat. Genet. 49, 451–456 (2017).

16. Uffmann, M. et al. Therapy reduction in patients with Down syndrome and myeloid leukemia: the international ML-DS 2006 trial. Blood 129, 3314–3321 (2017).

17. Laszig, S. et al. CPX-351 in Down syndrome-associated Myeloid Leukemia: Results and Prognostic Factors from the Phase 3 ML-DS 2018 Trial. Blood blood.2025030775 (2025) doi:10.1182/blood.2025030775.

18. Gruber, T. A. & Downing, J. R. The biology of pediatric acute megakaryoblastic leukemia. Blood 126, 943–949 (2015).

19. Suzuki, K. et al. A retrospective analysis of gene fusions and treatment outcomes in pediatric acute megakaryoblastic leukemia without Down syndrome. Haematologica 109, 1936–1940 (2024).

20. Athale, U. H. et al. Biology and outcome of childhood acute megakaryoblastic leukemia: a single institution’s experience. Blood 97, 3727–3732 (2001).

21. Llaurador, G. et al. Poor Outcomes Among Pediatric Non-Down Syndrome Acute Megakaryocytic Leukemia Patients Irrespective of Receipt of Hematopoietic Stem Cell Transplant. Blood 140, 10632–10633 (2022).

22. Schwarzer, A. et al. The non-coding RNA landscape of human hematopoiesis and leukemia. Nat. Commun. 8, 218 (2017).

23. Yeo, N. C. et al. An enhanced CRISPR repressor for targeted mammalian gene regulation. Nat. Methods 15, 611–616 (2018).

24. Filippova, J. A. et al. Are Small Nucleolar RNAs “CRISPRable”? A Report on Box C/D Small Nucleolar RNA Editing in Human Cells. Front. Pharmacol. 10, (2019).

25. DepMap 23Q4 Public. Figshare+ 10.25452/figshare.plus.24667905.v2 (2023).

26. Van Nostrand, E. L. et al. Robust transcriptome-wide discovery of RNA-binding protein binding sites with enhanced CLIP (eCLIP). Nat. Methods 13, 508–514 (2016).

27. Müller, S. et al. The oncofetal RNA-binding protein IGF2BP1 is a druggable, post-transcriptional super-enhancer of E2F-driven gene expression in cancer. Nucleic Acids Res. 48, 8576–8590 (2020).

28. Lemm, I. & Ross, J. Regulation of c-*myc* mRNA Decay by Translational Pausing in a Coding Region Instability Determinant. Mol. Cell. Biol. 22, 3959–3969 (2002).

29. Meristoudis, C. et al. Systematic analysis of the contribution of c-myc mRNA constituents upon cap and IRES mediated translation. Biol. Chem. 396, 1301–1313 (2015).

30. Sparanese, D. & Lee, C. H. CRD-BP shields c-myc and MDR-1 RNA from endonucleolytic attack by a mammalian endoribonuclease. Nucleic Acids Res. 35, 1209–1221 (2007).

31. Zhu, P. et al. A novel hypoxic long noncoding RNA KB-1980E6.3 maintains breast cancer stem cell stemness via interacting with IGF2BP1 to facilitate c-Myc mRNA stability. Oncogene 40, 1609–1627 (2021).

32. Zhou, H. et al. Regulatory mechanisms and therapeutic implications of insulin-like growth factor 2 mRNA-binding proteins, the emerging crucial m6A regulators of tumors. Theranostics 13, 4247–4265 (2023).

33. Hosono, Y. et al. Oncogenic Role of THOR, a Conserved Cancer/Testis Long Non-coding RNA. Cell 171, 1559–1572.e20 (2017).

34. Liu, S. et al. A novel lncRNA SNHG29 regulates EP300-related histone acetylation modification and inhibits FLT3-ITD AML development. Leukemia 37, 1421–1434 (2023).

35. Mahapatra, L., Andruska, N., Mao, C., Le, J. & Shapiro, D. J. A Novel IMP1 Inhibitor, BTYNB, Targets c-Myc and Inhibits Melanoma and Ovarian Cancer Cell Proliferation. Transl. Oncol. 10, 818–827 (2017).

36. Connerty, P. & Lock, R. B. The tip of the iceberg—The roles of long noncoding RNAs in acute myeloid leukemia. Wiley Interdiscip. Rev. RNA 14, e1796 (2023).

37. Luo, H. et al. *HOTTIP* lncRNA Promotes Hematopoietic Stem Cell Self-Renewal Leading to AML-like Disease in Mice. Cancer Cell 36, 645–659.e8 (2019).

38. Al-Kershi, S. et al. The stem cell–specific long noncoding RNA HOXA10-AS in the pathogenesis of KMT2A-rearranged leukemia. Blood Adv. 3, 4252–4263 (2019).

39. Stemmer, M., Thumberger, T., Keyer, M. del S., Wittbrodt, J. & Mateo, J. L. CCTop: An Intuitive, Flexible and Reliable CRISPR/Cas9 Target Prediction Tool. PLOS ONE 10, e0124633 (2015).

40. Adams, F. F. et al. An optimized lentiviral vector system for conditional RNAi and efficient cloning of microRNA embedded short hairpin RNA libraries. Biomaterials 139, 102–115 (2017).

41. Sanjana, N. E., Shalem, O. & Zhang, F. Improved vectors and genome-wide libraries for CRISPR screening. Nat. Methods 11, 783–784 (2014).

42. Mudge, J. M. et al. GENCODE 2025: reference gene annotation for human and mouse. Nucleic Acids Res. 53, D966–D975 (2025).

43. Volders, P.-J. et al. An update on LNCipedia: a database for annotated human lncRNA sequences. Nucleic Acids Res. 43, D174–D180 (2015).

44. Zhao, Y. et al. NONCODE 2016: an informative and valuable data source of long non-coding RNAs. Nucleic Acids Res. 44, D203–208 (2016).

45. Ng, M. et al. Myeloid leukemia vulnerabilities embedded in long noncoding RNA locus MYNRL15. iScience 26, (2023).

46. Li, W. et al. Quality control, modeling, and visualization of CRISPR screens with MAGeCK-VISPR. Genome Biol. 16, 281 (2015).

47. Cabianca, D. S. et al. A long ncRNA links copy number variation to a polycomb/trithorax epigenetic switch in FSHD muscular dystrophy. Cell 149, 819–831 (2012).

48. Bhayadia, R. et al. Endogenous Tumor Suppressor microRNA-193b: Therapeutic and Prognostic Value in Acute Myeloid Leukemia. J. Clin. Oncol. 36, 1007–1016 (2018).

49. Neumann, P. et al. The lncRNA GATA6-AS epigenetically regulates endothelial gene expression via interaction with LOXL2. Nat. Commun. 9, 237 (2018).

50. Cox, J. & Mann, M. MaxQuant enables high peptide identification rates, individualized p.p.b.-range mass accuracies and proteome-wide protein quantification. Nat. Biotechnol. 26, 1367– 1372 (2008).

51. Tyanova, S. et al. The Perseus computational platform for comprehensive analysis of (prote)omics data. Nat. Methods 13, 731–740 (2016).

52. The UniProt Consortium. UniProt: the Universal Protein Knowledgebase in 2025. Nucleic Acids Res. 53, D609–D617 (2025).

53. Dyer, S. C. et al. Ensembl 2025. Nucleic Acids Res. 53, D948–D957 (2025).

54. Quinlan, A. R. & Hall, I. M. BEDTools: a flexible suite of utilities for comparing genomic features. Bioinformatics 26, 841–842 (2010).

55. Harshil Patel et al. nf-core/rnaseq: nf-core/rnaseq v3.18.0 - Lithium Lynx. Zenodo 10.5281/ZENODO.1400710 (2024).

56. Ewels, P. A. et al. The nf-core framework for community-curated bioinformatics pipelines. Nat. Biotechnol. 38, 276–278 (2020).

57. Dobin, A. et al. STAR: ultrafast universal RNA-seq aligner. Bioinforma. Oxf. Engl. 29, 15–21 (2013).

58. Patro, R., Duggal, G., Love, M. I., Irizarry, R. A. & Kingsford, C. Salmon provides fast and bias-aware quantification of transcript expression. Nat. Methods 14, 417–419 (2017).

59. ComBat-seq: batch effect adjustment for RNA-seq count data | NAR Genomics and Bioinformatics | Oxford Academic. https://academic.oup.com/nargab/article/2/3/lqaa078/5909519.

60. Love, M. I., Huber, W. & Anders, S. Moderated estimation of fold change and dispersion for RNA-seq data with DESeq2. Genome Biol. 15, 550 (2014).

61. Zhu, A., Ibrahim, J. G. & Love, M. I. Heavy-tailed prior distributions for sequence count data: removing the noise and preserving large differences. Bioinforma. Oxf. Engl. 35, 2084–2092 (2019).

62. Liberzon, A. et al. The Molecular Signatures Database (MSigDB) hallmark gene set collection. Cell Syst. 1, 417–425 (2015).

63. Korotkevich, G. et al. Fast gene set enrichment analysis. 060012 Preprint at 10.1101/060012 (2021).

64. Ritchie, M. E. et al. limma powers differential expression analyses for RNA-sequencing and microarray studies. Nucleic Acids Res. 43, e47 (2015).

65. Hänzelmann, S., Castelo, R. & Guinney, J. GSVA: gene set variation analysis for microarray and RNA-seq data. BMC Bioinformatics 14, 7 (2013).

66. Brinkman, E. K., Chen, T., Amendola, M. & van Steensel, B. Easy quantitative assessment of genome editing by sequence trace decomposition. Nucleic Acids Res. 42, e168 (2014).

67. Luo, Y. et al. New developments on the Encyclopedia of DNA Elements (ENCODE) data portal. Nucleic Acids Res. 48, D882–D889 (2020).

