## Supplementary Information for "Lineage-restricted dependency on an oncofetal *SNHG29*-IGF2BP1 RNA axis in acute megakaryoblastic leukemia"

### Supplementary Figures

#### Supplementary Figure 1

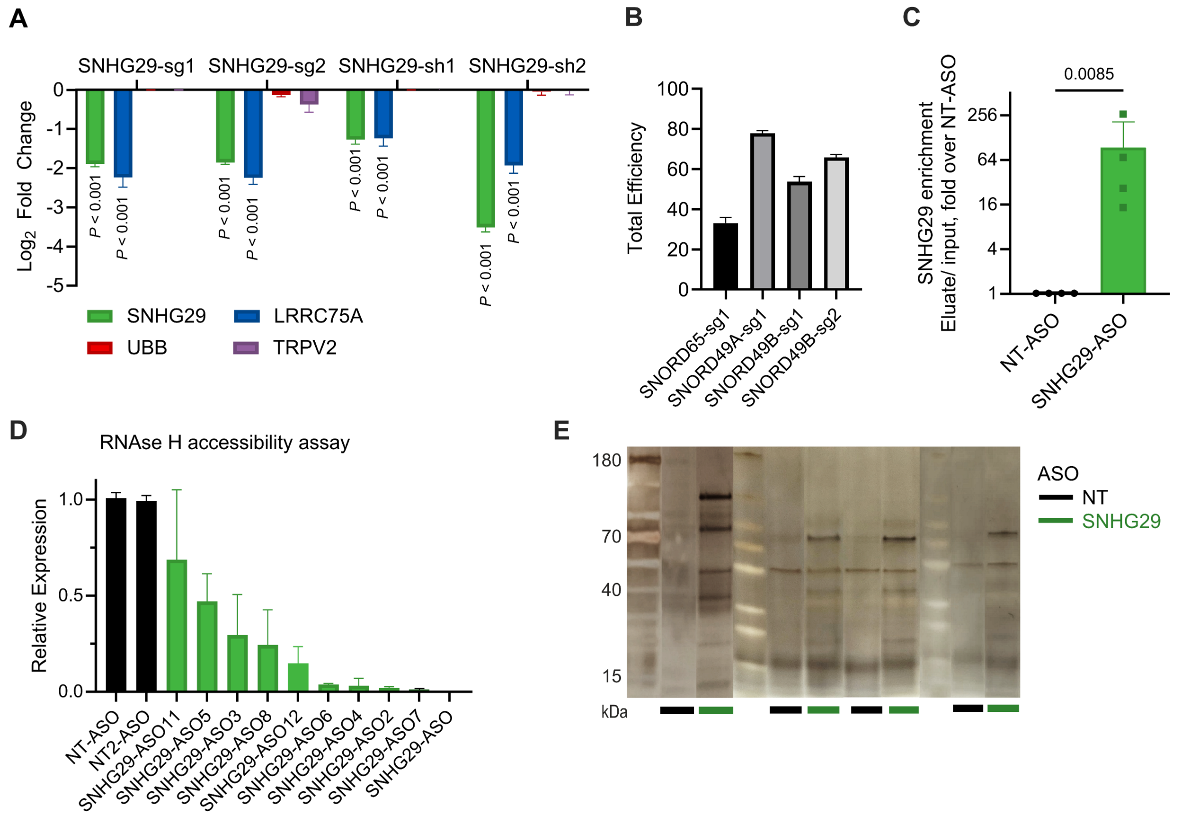


**Suppl. Fig. 1 |A** Expression changes of genes near the *SNHG29* locus in RNA sequencing following *SNHG29* knockdown in M-07e cells using CRISPRi with two different sgRNAs (left) or two different shRNAs (right) compared to the respective non-targeting control. Data shown are shrunken Log2 fold changes. ***FDR<0.001 (DESeq2). **B** CRISPR-Cas9 editing efficiency of indicated sgRNAs targeting SNORD65 and SNORD49A/B in M-07e Cas9 cells. Efficiency, represented as the percentage of total alleles with insertions/deletions (indels), was quantified by TIDE analysis following PCR amplification of the target sites (n=2, mean ± s.e.m.) **C** Relative *SNHG29* enrichment in the RNA affinity purification. Eluate was normalized to input and relative enrichment calculated over NT-ASO condition. (n=4 independent replicates; mean ± s.e.m.; Relative *SNHG29* enrichment compared using ratio paired t-test; **P<0.01) **D** RNase H accessibility assay using antisense DNA oligonucleotides (ASOs) tiling across the *SNHG29* transcript in M-07e cells. NT-ASO serves as a non-targeting control. Lower expression indicates higher accessibility of the target regions to RNase H-mediated degradation. Data are presented as relative expression normalized to NT-ASO control (n=2, mean ± s.e.m.). **E** Silver-stained SDS-PAGE gel of proteins eluted from the RNA pulldown assay. The gel displays proteins captured from cell lysates using either a biotinylated antisense oligonucleotide targeting *SNHG29* (ASO *SNHG29*) or a non-targeting control oligonucleotide (ASO NT). kDa: kDalton. (n=4 biological replicates).

**Supplementary Figure 2**

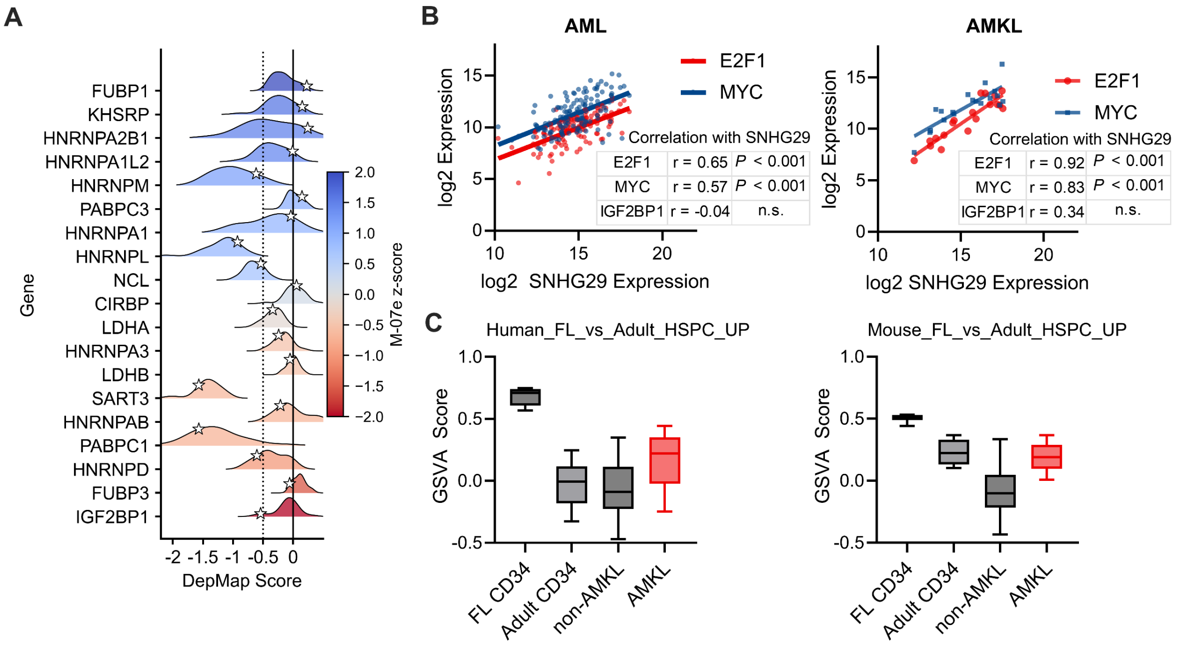


| AML | | |
|---|---|---|
| E2F1 | r = 0.65 | *P* < 0.001 |
| MYC | r = 0.57 | *P* < 0.001 |
| IGF2BP1 | r = -0.04 | n.s. |

| AMKL | | |
|---|---|---|
| E2F1 | r = 0.92 | *P* < 0.001 |
| MYC | r = 0.83 | *P* < 0.001 |
| IGF2BP1 | r = 0.34 | n.s. |

**Suppl. Fig. 2 | A** Dependency scores (DepMap 25Q3) of all proteins enriched in *SNHG29* pulldown. Specificity Z-scores were calculated across AML cell lines. **B** Correlation of E2F1, *MYC* and *IGF2BP1* (only statistical results shown) expression with *SNHG29* expression in non-megakaryoblastic AML (left, n=130) and AMKL (right, n=19) patient samples. Data are from RNA sequencing, Log2 transformed TPM. Pearson correlation co-efficients (r) and significance for E2F1, *MYC*, and *IGF2BP1* are indicated. ****P<0.0001; n.s., not significant. **C** Gene Set Variation Analysis (GSVA) scores of gene signatures upregulated in human (left) and mouse (right) fetal liver versus adult HSPCs in FL CD34+ (n=5), Adult CD34+ (n=8), non-AMKL (n=130) and AMKL (n=19) samples.

### Supplementary Tables

##### *Supplementary Table 1: CRISPRi lncRNA library*

##### *Supplementary Table 2: Clinical and genetic characteristics of patient cells used in this study*

| | Gender | Age (years) | WBC (x10³/µL) | Hemoglobin (g/dL) | BM blasts (%) | CNS | SCT | Molecular genetics | Cytogenetics | Response | Relapse |
|---|---|---|---|---|---|---|---|---|---|---|---|
| AMKL #1 | m | 1 | 40 | 4.59 | 64 | no | yes | KMT2A mutation | 46,XY[15]; FISH: negative (EVI1, RUNX1T1/RUNX1, MLL, CBFB, RARA) | CCR | yes |
| AMKL #2 | f | 3 | n/a | n/a | n/a | n/a | yes | CKIT mutation | 46,XX,add(3)(q?13), der(5) add(5)(p15) add(5)(q35) del(5) (q14q34),del(9) (q21q33), add(17) (q24)[4]/ 46,XX[11] | CCR | no |
| PICALMr | f | 5 | 111 | 9.5 | 93 | n/a | no | PICALM::MLLT10 | t(10;11)(p12;q14) | CCR | no |
| MLLr | f | 7 | 58.5 | 8.3 | 84 | no | yes | KMT2A::MLLT3 | 46,XX, t(9;11) (p22;q23) [8]/50, i-dem, +3, +8, +18, +19[15] | CCR | no |

##### *Supplementary Table 3:* sgRNA and shRNA name spacer sequences

| Oligonucleotide name | System | Sequence |
|---|---|---|
| LUC-sg | CRISPRi | AAGAGATACGCCCTGGTTCCTGG |
| SNHG29-sg1 | CRISPRi | GGTAGGCTCTCTAGGAATCTGGG |
| SNHG29-sg2 | CRISPRi | AAGCCTGTCTAGAGGGCGGAGGG |
| SNHG29-sg3 | CRISPRi | GTTATGGAGGTAGGCTCTCTAGG |
| Ctrl-sh | shRNA | AGGAATTATAATGCTTATCTA |
| SNHG29-sh1 | shRNA | GAATCAGCATCATGTTTGGCA |
| SNHG29-sh2 | shRNA | GAAAAGGCACATTGGGTATCA |
| LUC-Rx | CRISPR-CasRx | TGCGTCGGTAAAGGCGATGGTG |
| SNHG29-Rx1 | CRISPR-CasRx | CCTCTGATACATAAGGCAAGCAT |
| SNHG29-Rx2 | CRISPR-CasRx | CCAGCTCTAAAACAGCACTCTGT |
| SNHG29-Rx3 | CRISPR-CasRx | GGCTCCAATACTCAGCTGCCAAA |
| LRRC75A-sg1 | CRISPR-Cas9 | ATACAGAACGTCGTCTAGCG |
| LRRC75A-sg2 | CRISPR-Cas9 | AGGTCAGTGGGATTCCCGAC |
| LRRC75A-sg3 | CRISPR-Cas9 | TCAATGGCAACCGGTTGACC |
| LRRC75A-sg4 | CRISPR-Cas9 | ATACAGAACGTCGTCTAGCG |
| SNORD65-sg1 | CRISPR-Cas9 | TTCACCACTACACAATCTGCCGG |
| SNORD49A-sg1 | CRISPR-Cas9 | AGACTTGACTGCAATCAGACAGG |

| | | |
|---|---|---|
| SNORD49B-sg1 | CRISPR-Cas9 | TTCCTATTACAAGTATCATCAGG |
| SNORD49B-sg2 | CRISPR-Cas9 | GTCCTGATGATACTTGTAATAGG |
| IGF2BP1-sg1 | CRISPR-Cas9 | CAAGATCATCTTACAAGCGG |
| IGF2BP1-sg2 | CRISPR-Cas9 | AATGGCACCCACATACTGGG |
| IGF2BP1-sg3 | CRISPR-Cas9 | CTCGTCCGGGCAGTCCACGA |

#### *Supplementary Table 4: PCR primers*

| Target | Application | Direction | Sequence |
|---|---|---|---|
| SNHG29 | qRT-PCR | Forward | CATGGTCCAGGAGCTGCTG |
| | | Reverse | AGCGATACAGAACGTCGTCTAG |
| B2M | qRT-PCR | Forward | TCTCTCTTTCTGGCCTGGAG |
| | | Reverse | AATGTCGGATGGATGAAACC |
| SNORD49A & B | TIDE PCR | Forward | GTATTGGACAGCCTGGCAGG |
| | | Reverse | CCAGCCTCAGGGGAGTTGTA |
| SNORD65 | TIDE PCR | Forward | GCTGTTTTAGAGCTGGCAGC |
| | | Reverse | GTGGGAGGATTGCTTAGGCC |

#### *Supplementary Table 5: Antisense oligos*

| ASO name | Application | Sequence |
|---|---|---|
| SNHG29-ASO | RNA pulldown | mGmUmAmAmUmGmAmAmUmG-<br>mAmUmAmCmCmCmAmAmUmGmU<br>/iSp9/rArCrGrArUrC/3deSBioTEG/ |
| NT-ASO | RNA pulldown | mGmGmAmCmGmAmUmUmCmG-<br>mAmUmCmGmAmUmAmAmUmCmU<br>/iSp9/rArCrGrArUrC/3deSBioTEG/ |
| NT-ASO | Accessibility assay | GGACGATTCGATCGATAATCT |
| NT2-ASO | Accessibility assay | GCAAGGAACGTGTGAGACTA |
| SNHG29-ASO2 | Accessibility assay | ATTCTTCACGAATTTGCAACC |
| SNHG29-ASO3 | Accessibility assay | AATACTCAGCTGCCAAACATG |
| SNHG29-ASO4 | Accessibility assay | TTCTCAAAACCTCATGGCAGG |
| SNHG29-ASO5 | Accessibility assay | CAGCTCTAAAACAGCACTCTG |
| SNHG29-ASO6 | Accessibility assay | GGAATATAACCTTCTCTTGGG |
| SNHG29-ASO7 | Accessibility assay | AATCCAAACTGATGGCAGCTA |
| SNHG29-ASO8 | Accessibility assay | TTGATGCCAGTTAGTTTTCAG |
| SNHG29-ASO11 | Accessibility assay | ATATATCTCTTGGATCTGCTG |
| SNHG29-ASO12 | Accessibility assay | CCTTTTCTAGAAAAAGTTGCC |
| SNHG29-ASO | Accessibility assay | GTAATGAATGATACCCAATGT |
